# Theobromine is Associated with Slower Epigenetic Ageing

**DOI:** 10.1101/2025.04.15.648884

**Authors:** Ramy Saad, Ricardo Costeira, Pamela R Matias Garcia, Christian Gieger, Karsten Suhre, Annette Peters, Gabi Kastenmüller, Ana Rodriguez-Mateos, Cristina Dias, Cristina Menni, Melanie Waldenberger, Jordana T Bell

## Abstract

Theobromine, a commonly consumed dietary alkaloid derived from cocoa, has been linked to extended lifespan in model organisms and to health benefits in humans. We examined associations between circulating theobromine intake, measured using serum metabolomics, and blood-based epigenetic markers of biological ageing in two European human population-based cohorts. Serum theobromine levels were significantly associated with reduced epigenetic age acceleration, as measured by GrimAge (p<2e-7) and DNAmTL (p<0.001) in over 500 individuals from the TwinsUK cohort, and both signals replicated in 1,160 individuals from the KORA cohort (p = 7.2e-08 and p = 0.007, respectively). Sensitivity analyses including covariates of other cocoa and coffee metabolites suggest that the effect is specific to theobromine. Our findings indicate that the reported beneficial links between theobromine intake on health and ageing extend to the molecular epigenetic level in humans.

## Introduction

Dietary phytochemicals are compounds found in plants that have been reported to benefit human health. They include polyphenols, alkaloids, terpines, flavonoids, and others (1). Evidence from both epidemiological and human intervention trials have identified beneficial effects of various phytochemicals on health and ageing, including on biomarkers of cholesterol transport (2), inflammation (3), and cellular senescence (4).

Alkaloids in plants form a large component of dietary phytochemicals, as they are both abundant and highly bioactive (5). This bioactivity is a function of their purpose as protective chemicals and, therefore, alkaloids have wide-ranging *in vivo* actions, along with narrow therapeutic indices. Specifically, alkaloids have been studied for their relevance to age-related diseases, including cancers (6), type 2 diabetes (7) and inflammation (8). Notable examples of pharmacologically active alkaloids include indole (9), indolizidine (10), as well as specific subtypes such as berberine (11), morphine, strychnine, quinine, and others (5).

Coffee and cocoa are widely consumed foods, associated with reduced cardiovascular disease (CVD) and mortality (12, 13). Cocoa and coffee share several important alkaloids including the methylxanthines theobromine (TB), caffeine (CAF), theophylline (TP), paraxanthine (PX) and 7-methylxanthine (MX) (14) *(Figure 1a).* The coffee-associated methylxanthines (CAF, TP and PX) are found in lower concentrations in cocoa (8). TB and MX, are partial metabolites of CAF, though both are also found in much higher concentrations in cocoa as primary unprocessed metabolites (15). TB has previously been linked to multiple aspects of health and ageing. For example, studies in model organisms have identified links between TB and extended lifespan (16). Furthermore, multiple observational human cohort studies have reported clear links between TB intake and various aspects of improved health (17). Despite this, the exact impacts of TB on health and ageing are still not fully understood, and the molecular pathways that underlie these effects are largely unknown.

**Figure 1.**
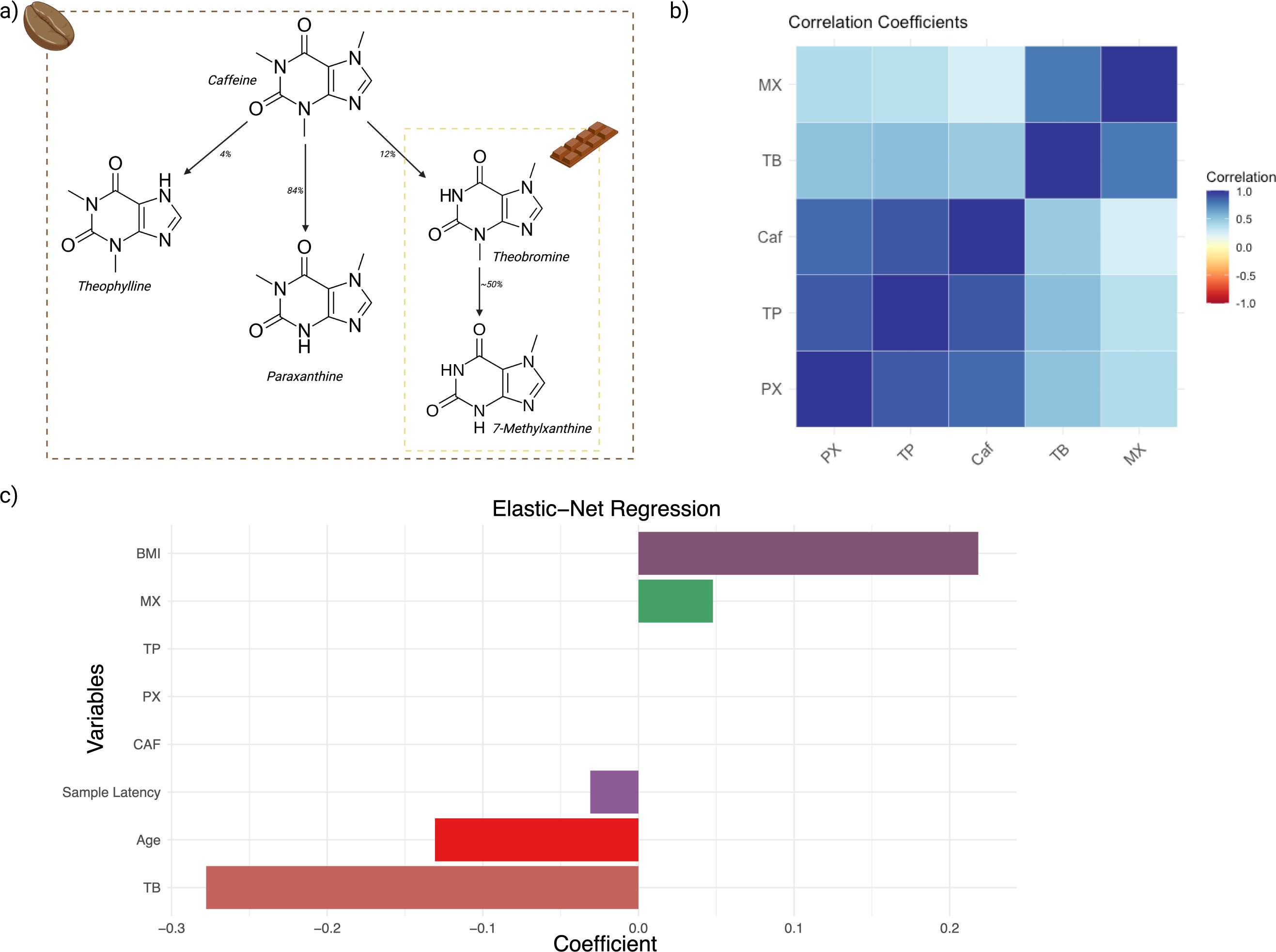
Major dietary sources of methylxanthines and their correlations in the TwinsUK sample. 1a) Schematic presenting key methylxanthines, their respective dietary sources and their derivation as secondary metabolites. 1b) Correlation heatmap of coffee-related metabolites in the TwinsUK sample. 1c) Bar plot representation of the Elastic-net Regression coefficients with 10-fold cross-validation of variables against GrimAgeAccel in the TwinsUK sample.

Multiple biological mechanisms can mediate the effects of dietary phytochemicals on human health and ageing, and one of these is the epigenetic regulation of gene expression. Alkaloids can influence epigenetic processes, for example, through inhibition of histone deacetylases or DNA methyltransferases (DNMTs) (18).

Cocoa and coffee consumption have been linked to multiple DNA methylation changes in humans, where extracts from cocoa can affect global leukocyte DNA methylation levels potentially though inhibition of DNMTs (19), and distinct blood DNA methylation signals have been associated with coffee consumption (20). Therefore, alkaloids, such as those found in cocoa, may exert their beneficial effects on health and ageing potentially through changing the human epigenome.

Epigenetic deregulation is a key hallmark of ageing. The effects of ageing on genome-wide methylation have been widely documented, including reduction of global DNA methylation (21), global increase in Shannon entropy of methylation patterns (22), and site-specific changes in differential and variable DNA methylation levels (23–25). Hence, multiple studies have developed epigenetic clocks towards predicting different age-related features, such as chronological age (26), time to death (27), pace of ageing (28), as well as other molecular biomarkers of ageing including telomere length (29). As such, epigenetic clocks may act as useful tools for assessing whether specific dietary phytochemicals are associated not only with epigenetic modifications, but also with the rate of ageing, as measured by these clocks.

Several recent studies have investigated the association of nutrients and metabolites to epigenetic ageing. Early studies focused on diet questionnaire data, identifying compelling associations between B vitamin intakes and with slower epigenetic ageing (30). Smaller-scale intervention trials have also explored the impact of dietary changes on epigenetic age. For example, an 8-week randomised controlled trial intervention in six post-menopausal women found that an increase in dietary polyphenols resulted in significant deceleration of epigenetic aging, as measured by the Horvath clock (31). Moreover, dietary interventions such as calorie restriction can also influence epigenetic aging, as identified from the CALERIE trial using the DunedinPACE epigenetic clock (32). However, results were not consistent across different epigenetic clocks, highlighting potential variability in how they capture ageing processes.

In this study, including two independent human population-based cohorts, we investigated whether individual bioactive alkaloids in coffee and cocoa are associated with reduced epigenetic ageing, and may therefore potentially contribute towards extension of human healthspan.

## Results

We initially tested for the association between six metabolites found in coffee and cocoa, and epigenetic measures of ageing in blood samples from 509 healthy females from the TwinsUK cohort (median age = 59.8, IQR = 12.81, BMI = 25.35). The six metabolites included the methylxanthines CAF, TP, TB, MX and PX and theanine, and biological ageing analyses focused on GrimAgeAccel. TB was significantly associated with reduced epigenetic ageing as captured by GrimAgeAccel (B = -1.576, standard error = 0.3, p = 3.99e-6) *(Figure 2, Supplementary Table 1).* Extending analyses to test for association between TB and other measures of biological ageing, including methylation markers of telomere length, PhenoAge and DunedinPACE, we identified another significant association with DNAmTL (B = 0.03, standard error = 0.0124, p = 0.0029) *(Figure 2, Supplementary Table 1)*.

**Figure 2.**
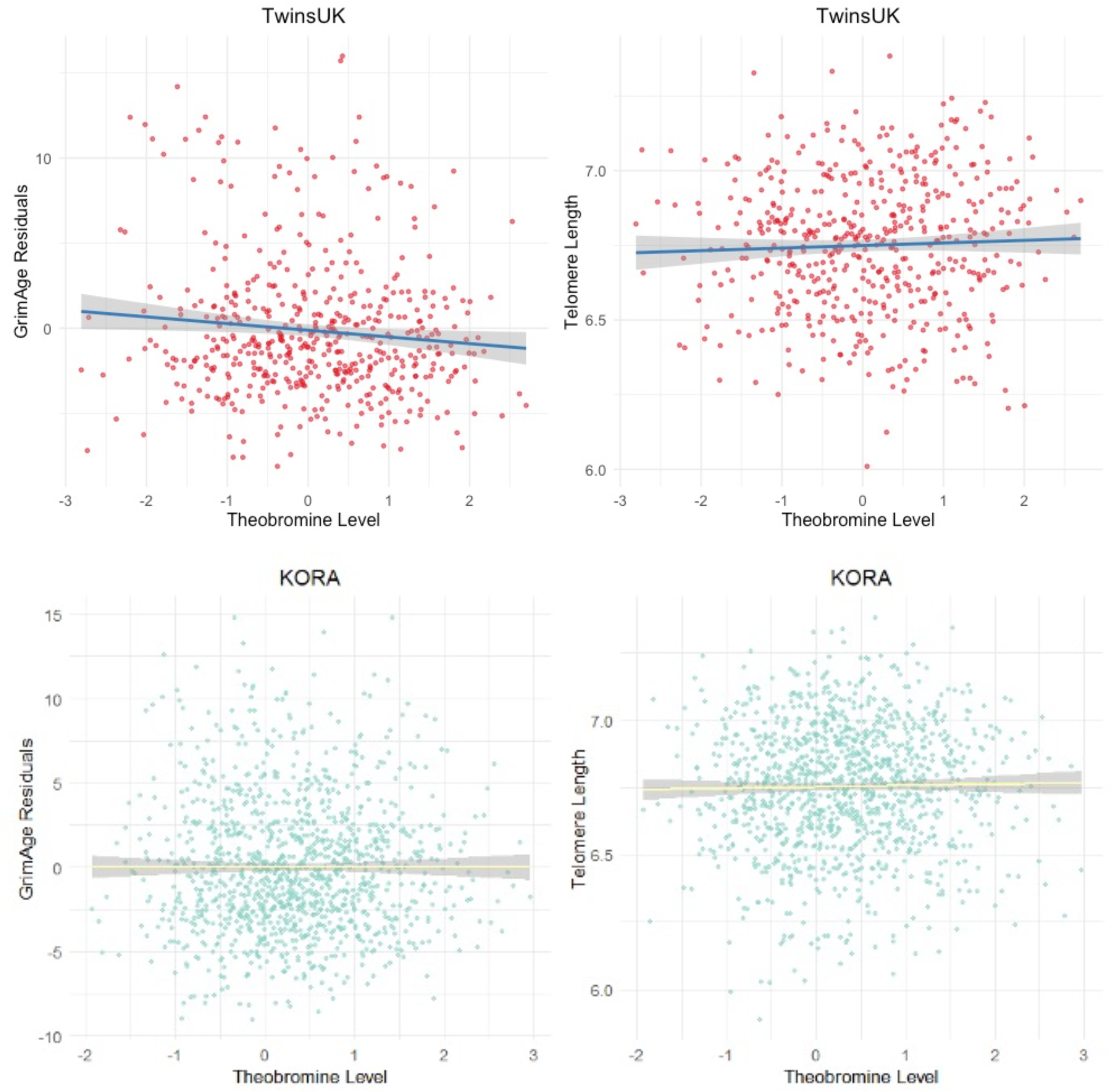
The association between TB and epigenetic age in the TwinsUK cohort. Scatter plots of the GrimAge acceleration residuals (left) and epigenetic estimate of telomere length (DNAmTL, right) in the discovery TwinsUK sample of 509 females (top row) and KORA sample of 1160 individuals (bottom row). Lines of best fit and standard errors are labelled.

As both cocoa and coffee include TB, we carried out further analyses to dissect the correlations between methylxanthines and food component intakes within the TwinsUK sample. Correlation coefficients support the expected patterns of close correlation among coffee-associated methylxanthines (CAF, TP, PX) and close correlation among the cocoa-associated methylxanthines (TB and MX) (*Figure 1b*), demonstrating consistency with the undertaken metabolomic analysis.

Specifically, we observed that coffee-associated methylxanthines CAF and TP were strongly correlated to each other (R=0.89), and that cocoa-associated methylxanthines TB and MX were also strongly correlated to each other (R=0.78). In contrast, TB and CAF showed only moderate correlation (R=0.46) *(Figure 1b)*. The weaker correlation between TB and CAF in our cohort reflects the expected low metabolism of CAF to TB, and the likely differential food sources of these metabolites *in vivo* (33). Indeed, TB was previously associated with chocolate consumption in a larger sample from the same TwinsUK population cohort (B = 0.024, p = 1.34e-11) (34). In the current sample (509 twins), we confirm a positive, but weaker correlation between the consumption of ‘chocolate’ (as reported by food frequency questionnaires) and TB levels (R = 0.136). TB consumption was not strongly associated to diet quality (AHEI, R = -0.0293).

Several sensitivity analyses were undertaken to mitigate the effects of potential dietary intake confounders. First, we re-examined the association between TB and GrimAge acceleration, but now including CAF, TP, PX, and MX as additional covariates. We then carried similar analyses, also including TP, PX, and MX as additional covariates because they are metabolite derivatives of CAF (14) (*Figure 1a)*. The associations between TB and slower epigenetic ageing remained significant in these extended models with additional covariates *(Supplementary Figure 1a,b; Supplementary Table 1)*, suggesting that effects are specific to TB and not an alternative xanthine derivative. Furthermore, we re-analysed the data to assess the effect of time latency between date of DNA methylation and metabolomic sample collection. Samples were subset by window of latency periods, including 5 years (n = 509), 2 years (n = 420), 1 year (n = 276) and contemporaneous (same-day) sampling (n = 121). The strength of the association increased with shorter latency periods *(Supplementary Figure 1c)*.

Targeted replication of the TB and epigenetic ageing rate associations was sought in a larger sample of 1,160 individuals (median age = 60, median BMI = 27) from the KORA cohort (35), where serum metabolomic and DNA methylation profiles were obtained from the same time-point. We replicated the association between reduced GrimAgeAccel and TB in a model including all technical and biological covariates of our study (CAF, TP, PX, MX; coefficient = -1.06, standard error = 0.195, p = 7.177E-08) *(Figure 2)*. We also replicated the association of TB with DNAmTL (coefficient = 0.022, standard error = 0.008, p=0.007) *(Figure 2)*.

As a further follow up analysis, we also stratified the TwinsUK sample by smoking status. The reduced epigenetic ageing acceleration signal was most significant in previous and current smokers (B = -2.687, p = <2.2e-16, n = 53), compared to never smokers (*Figures 3a, 3b)*. Nicotine likely induces the breakdown of TB by enzyme induction (36) and may influence the pharmacodynamic clearance or bioavailability of TB and its byproducts. Alternatively, the ageing-protective effects of TB could be more pronounced in smokers. Further research is needed in model organisms to confirm these differential effects.

A final set of follow-up analyses explored feature selection using LASSO and elastic net regression to assess which metabolites most strongly relate to the epigenetic measures of biological ageing. LASSO regression with GrimAgeAccel as the dependent variable and all technical covariates and metabolites as independent variables identified TB as a significant predictor for GrimAgeAccel (coefficient = -0.231; RMSE = 3.644), with similar results using 10-fold cross-validation (coefficient = -0.186; RMSE = 3.834). Elastic-net regression with 10-fold cross-validation (best alpha: 0.2, best lambda: 0.419) showed consistent results (GrimAgeAccel TB coefficient = -0.277; RMSE = 3.8) (*Figure 1c)*.

## Discussion

Here we report a significant association between circulating levels of theobromine (TB) with slower epigenetic ageing in two independent population-based cohorts. TB is a relatively unexplored dietary phytonutrient that has recently been linked to beneficial health effects and extended lifespan in model organisms (16). However, there have been limited studies of the role of TB in human cohorts.

The association of TB and biological ageing measures is most pronounced by the GrimAge epigenetic clock acceleration measures, which strongly predicts time to death. The pattern was also captured by DNAmTL, which estimates telomere length. The two epigenetic ageing measures, DNAmTL and GrimAge acceleration residuals are weakly correlated (R = 0.29) in the TwinsUK sample, and this supports previous reports in the literature that telomere length and genome-wide epigenetic ageing are independently associated with ageing (37). Previous work has explored how epigenetic clocks may capture different mechanisms underpinning hallmarks of ageing, such as telomere attrition and epigenetic ageing (38). We therefore considered the two measures to capture separate aspects of the ageing process, that do not necessarily overlap.

Methylxanthines are found across various food groups in different proportions, with CAF being the most prominent in coffee, and TB being the most prominent in cocoa (39). Exact proportions can vary across foods and also depend on food quality, processing methods (such as decaffeination), or inter-individual variability (such as genetic variation in monooxygenase function or presence of exogenous P450 enzyme inducers or inhibitors (40)). Our sensitivity analyses support the conclusion that the association effect is specific to TB and is likely not attributed to CAF, TP, PX or MX. This conclusion stems from results based on accounting for multiple metabolites as covariates in the linear association models, and results from 10-fold cross-validation LASSO and elastic-net regression. This suggests that TB may affect a common biological pathway relevant to ageing.

Several studies predominantly in model organisms have identified links between TB and improved aspects of health and ageing. Importantly, TB has been reported to extend lifespan in ROS-sensitive strains of *C. elegans* (16). It has also been noted to have differential psychotropic actions to caffeine (41). In mice, modest supplementation of 0.05% TB results in significant increases in the neurotrophic factors CREB and BDNF, which are relevant to reward and learning (42), but higher doses of TB were associated with better lipid profiles and lower blood pressure in a retrospective cross-sectional study (43). Although some methylxanthines are used in clinical practice (44), TB has not been explored in depth for its medical utility, but it has been suspected to be of importance to human health (45).TB has also been previously associated with the enrichment of beneficial microbiota with SFCA-producing abilities (46). Future studies should explore if the gut microbiome composition may mediate the effect of TB on human health and ageing.

One plausible explanation for the correlation between TB and epigenetic age is whether it may be a biomarker for a collinear confounder. For instance, TB may signify flavan-3-ol consumption, as these (poly)phenols are abundant in cocoa but were not available in the metabolomic data. Methylxanthines, including theobromine, have been shown to enhance the vascular effects of flavan-3-ols by improving endothelial function and increasing nitric oxide bioavailability; however, when administered alone, they did not elicit any effect (47) and the cardiometabolic and healthy aging benefits of flavan-3-ols are well established (13, 48). On the other hand, the sensitivity analysis using elastic-net regression supports the conclusion that the effect is specific TB and not other collinear methylxanthines, making the possibility of a hidden confounding variable less likely. Further research is needed to disentangle the potential mechanisms by which TB is associated with reduced epigenetic ageing and exclude any potential confounders not assessed by our study or reverse-causality.

In conclusion, our study identifies strong links between TB and measures of epigenetic ageing, suggesting that TB is relevant to human ageing. Further exploration of TB and age-related health markers may identify key epigenetic mechanisms transducing this effect and reveal a potential use of TB towards extending the human healthspan.

## Materials and Methods

### Discovery Cohort Data

The discovery sample in this study included 509 monozygotic and dizygotic twin female participants from the TwinsUK cohort (49). Metabolomic data in these participants were generated in fasting serum samples using the Metabolon Inc mass spectrometry platform (Metabolon, Inc., Durham, NC). Metabolite concentrations were measured at fasting from serum, samples by Metabolon Inc. (Durham, USA) using an untargeted Liquid chromatography–mass spectrometry (LC-MS) platform as previously described (50). Metabolites with more than 20% missingness and metabolite data outliers (±3SD from mean) were excluded. The remaining metabolites were day median-normalised, imputed to the day minimum, and inverse-normalised. Six metabolites associated with coffee or cocoa consumption were analysed in this study, including theobromine (TB), caffeine (CAF), theophylline (TP), paraxanthine (PX), 7-methylxanthine (MX) and theanine, an amino acid prevalent in tea.

Dietary intakes in the TwinsUK sample were estimated using a modified version of the European Prospective Investigation into Cancer and Nutrition (EPIC) food frequency questionnaire (FFQ). This version incorporates food items from the EPIC Norfolk study (51). FFQs were excluded if more than 10 food items were unanswered or if the total energy intake, derived from the FFQ and expressed as a ratio of the subject’s estimated basal metabolic rate (calculated using the Harris– Benedict equation), fell outside 2 standard deviations of the mean (below 0.52 or above 2.58), as previously described (52).

Whole blood DNA methylation profiles were generated for the same 509 participants in the discovery TwinsUK sample using the Infinium HumanMethylation450 BeadChip (Illumina). Epigenetic data generation and processing has previously been described (53, 54). Briefly, minfi (55) was used to exclude samples with median methylated/unmethylated ratio < 10.5, and Enmix (56) was used for background correction, dye bias correction and quantile normalisation of the data. Methylation beta-values were estimated for signals with detP < 0.000001 and nbead > 3. Finally, probes and samples with >5% missingness were excluded, as were any outlier samples identified by Enmix (56). Polymorphic or cross-reactive probes were removed. Mass spectrometry and DNA methylation data were not always obtained from the same clinical visit and samples were selected at a maximum of 5 years apart (median = 0.11 years). Blood cell proportions were estimated following the Houseman et al. method (57) and obtained from Horvath’s calculator.

### Epigenetic clocks

Epigenetic clocks in this study were estimated using Horvath’s ’New Methylation Age Calculator’. The analyses focused on epigenetic age acceleration estimated as the residuals of epigenetic age adjusted for chronological age as estimated in Horvath’s calculator. Analyses focused on two epigenetic clocks including GrimAge acceleration (GrimAgeAccel), selected due to its high predictive ability for time to death (27) and previous use in similar work related to diet quality (58); and a DNA methylation-based estimator of telomere length (DNAmTL) (29). Additional analyses extended to other epigenetic clocks including the Hannum clock (25), PhenoAge (59), and DunedinPACE (28).

### Replication Cohort Data

Replication was undertaken in 1,160 fasting serum samples from the KORA (Cooperative Health Research in the Region of Augsburg) cohort. The KORA (Kooperative Gesundheitsforschung in der Region Augsburg) F4 (2006–2008) is a follow-up study from the KORA S4 (n=4,261) survey carried out 1999-2000 (35). Fasting blood serum samples were collected from participants of the KORA study population and profiled using the Metabolon platform (Metabolon, Inc., Durham, NC), as described previously (60). Median-normalisation was achieved by multiplying each metabolite with overall median values and log-transformed. Whole blood DNAm profiles in the KORA cohort were generated using the HumanMethylation450 BeadChip, and processing of data has previously been described (61). Estimation of epigenetic ageing clocks followed the methodology outlined for the discovery sample analysis.

### Statistical Analysis

Association analyses were carried out in RStudio (2023.09.1+494) using linear mixed-effects models (R package ‘lme4’). Epigenetic acceleration measures were the response variable and theobromine levels were the predictors. Models were adjusted for covariates including blood cell type proportions, age and body-mass index (BMI) as fixed-effect variables, and for family relatedness as a random effect term. The primary analysis investigated associations between each metabolite and GrimAgeAccel and DNAmTL. Extended analyses also considered additional epigenetic clocks (Hannum clock, PhenoAge, and DunedinPACE).

Sensitivity analyses included additional covariates CAF, TP, PX and MX to account for potential confounding across food components. As multicollinearity is a potential confounder, two additional sensitivity analyses were undertaken, 10-fold cross validated LASSO and elastic-net regression (R package ‘glmnet’). LASSO regression penalises coefficients by shrinkage, reducing the impact of individual multicollinear variables. Elastic-net regression also uses penalisation to regularise results and reduce the influence of collinear metabolites, by using a composite of LASSO and Ridge regularisation methods to enable best fit.

## Abbreviations

(CAF): Caffeine
(DNAm): DNA methylation
(MX): 7-methylxanthine
(PX): Paraxanthine
(TB): Theobromin
(TP): Theophylline

## Author Contributions

RS and JTB conceived the study. RS and RC carried out the primary analysis, and PRMG contributed to the analysis. CG, KS, AP, GK, CM, and MW contributed reagents, materials and input on results interpretation. ARM and CD contributed to results interpretation and write-up. RS, RC, and JTB wrote the manuscript. All authors reviewed and approved the manuscript.

## Acknowledgements

The authors thank all research volunteers who participated in the TwinsUK and KORA studies. In addition, we thank all participants for their long-term commitment to the TwinsUK and KORA studies, the staff for data collection and research data management, and the members of the TwinsUK Resource Executive Committee and of the KORA Study Group (https://www.helmholtz-munich.de/en/epi/cohort/kora) who are responsible for the design and conduct of the TwinsUK and KORA studies, respectively.

The authors also acknowledge use of the research computing facility at King’s College London, the King’s Computational Research, Engineering and Technology Environment (CREATE) (https://doi.org/10.18742/rnvf-m076).

## Conflict of Interest

Nothing to declare.

## Ethical statement

Ethical approval for the discovery sample analyses in TwinsUK was granted under TwinsUK BioBank ethics, approved by North West – Liverpool Central Research Ethics Committee (REC reference 19/NW/0187). The KORA study was performed in line with the principles of the Declaration of Helsinki. Study methods were approved by the Ethics Committee of the Bavarian Chamber of Physicians (REC: #06068).

## Consent

Written informed consent was obtained from all research participants prior to taking part in any research activities.

## Funding

This project was supported by the European HDHL Joint Programming Initiative funding scheme DIMENSION project (BBSRC BB/S020845/1 and BB/T019980/1 to J.T.B.). The TwinsUK study is funded by the Wellcome Trust, Medical Research Council, Versus Arthritis, European Union Horizon 2020, Chronic Disease Research Foundation (CDRF), ZOE LIMITED, and the National Institute for Health Research (NIHR) Clinical Research Network (CRN) and Biomedical Research Centre based at Guy’s and St Thomas’ NHS Foundation Trust in partnership with King’s College London. The KORA study was initiated and financed by the Helmholtz Zentrum München – German Research Center for Environmental Health, which is funded by the German Federal Ministry of Education and Research (BMBF) and by the State of Bavaria. Data collection in the KORA study is done in cooperation with the University Hospital of Augsburg. The project DIMENSION, partnering site Helmholtz Munich, received financial support by a grant of the European HDHL Joint Programming Initiative funding scheme, administered by the Federal Ministry of Research in Germany, Grant No. 01EA1902A (M.W.). K.S. is supported by the Biomedical Research Program at Weill Cornell Medicine in Qatar, a program funded by the Qatar Foundation and by Qatar National Research Fund (QNRF) grants NPRP11C-0115-180010 and ARG01-0420-230007.

---

Supplementary Figure 1. Sensitivity analyses exploring the specificity of the theobromine association with epigenetic ageing measures in the TwinsUK sample. 3a) and 3b) Theobromine and GrimAge association significance in never smokers and current/ex-smokers. Covariates in the models included ‘N’ (BMI + Age + Cell Proportions), as well as ‘N’ and methylxathines CAF, TP, PX and MX, as labelled on the Figure. TB coefficient effect sizes are shown. 3c) Magnitude of effect sizes of DNAmTL association with GrimAgeAccel. As the scales and direction differ (with negative GrimAgeAcel and positive DNAmTL both indicating a slower age) the regression coefficient values of GrimAgeAccel were converted to absolute values and maximum values of DNAmTL were scaled to GrimAgeAccel (scale factor: 43.43). Correlation lines between latency values are applied (linear model) and standard errors are shown.

Supplementary Table 1. Association results in TwinsUK cohort. Sample Latency is provided in years between methylation and metabolomic measurements. Covariates included in the linear model are ‘N’ (BMI + Age + Cell Proportions) and methylxathines CAF, TP, PX and MX as labelled.

## Notes

### Competing Interest Statement

The authors have declared no competing interest.

