## Supplementary figures and images for "Theobromine is Associated with Slower Epigenetic Ageing"

### Supplemental Figure

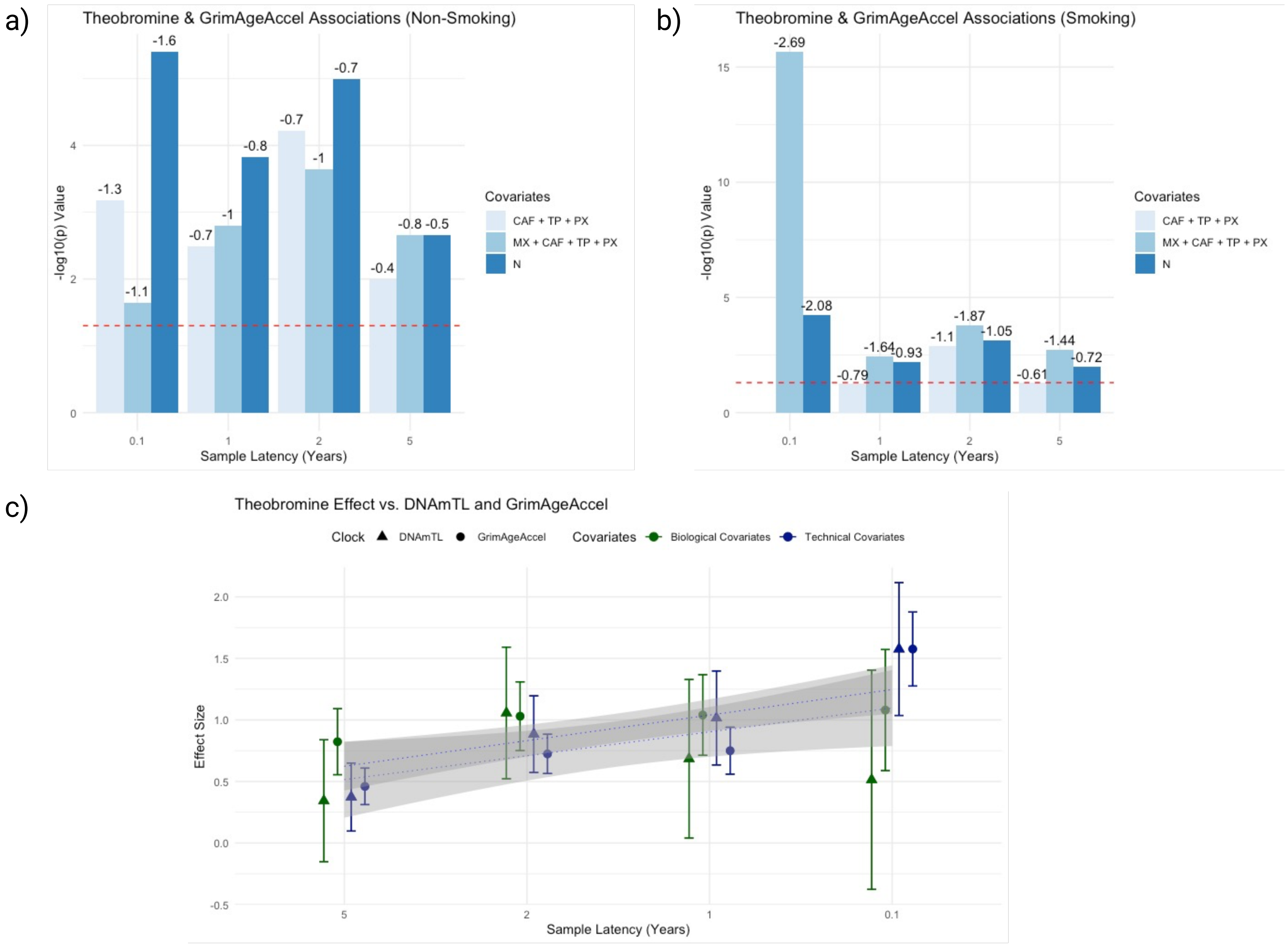
