## Supplemental Table for "Theobromine is Associated with Slower Epigenetic Ageing"

| Sample Latency (yrs) | n | Covariates | GrimAge | | | DNAmTL | | | DunedinPACE | | | PhenoAge | | | AgeAccelHannum | | |
|---|---|---|---|---|---|---|---|---|---|---|---|---|---|---|---|---|---|
| | | | *β* | *se* | *p* | *β* | *se* | *p* | *β* | *se* | *p* | *β* | *se* | *p* | *β* | *se* | *p* |
| 5 | 509 | N | -0.46 | 0.148352 | 0.002227 | 0.008597492 | 0.006367478 | 0.1784 | 0.002433033 | 0.003743138 | 0.5135 | -0.1474319 | 0.2116257 | 0.4955 | 0.2210168 | 0.179496 | 0.2135 |
| 2 | 420 | N | -0.724 | 0.1596351 | 1.03E-05 | 0.020360325 | 0.007167899 | 0.004462 | -0.0068576 | 0.004158709 | 0.09612 | -0.3780389 | 0.245318 | 0.128 | 0.03520191 | 0.20896333 | 0.8581 |
| 1 | 276 | N | -0.75 | 0.1907382 | 0.00015 | 0.02338561 | 0.00878681 | 0.00798 | -0.004733561 | 0.004859301 | 0.3249 | -0.4952221 | 0.3012944 | 0.09743 | -0.06744427 | 0.24238152 | 0.7961 |
| 0.1 | 121 | N | -1.576 | 0.3005898 | 3.99E-06 | 0.03628296 | 0.01242632 | 0.002985 | -0.007792432 | 0.006966381 | 0.3101 | -0.3016093 | 0.4554014 | 0.5374 | -0.04750247 | 0.33094191 | 0.918 |
| 5 | 509 | CAF + TP + PX | -0.44 | 0.1698753 | 0.01004 | 0.010702108 | 0.007202701 | 0.1362 | -0.001484194 | 0.004284681 | 0.7303 | -0.1722435 | 0.2427985 | 0.4777 | 0.2069323 | 0.205579 | 0.3087 |
| 2 | 420 | CAF + TP + PX | -0.734 | 0.1795713 | 6.07E-05 | 0.021952714 | 0.007920683 | 0.005354 | -0.006831862 | 0.004684106 | 0.1399 | -0.366313 | 0.2735349 | 0.1787 | 0.02361039 | 0.23384471 | 0.9211 |
| 1 | 276 | CAF + TP + PX | -0.667 | 0.2215733 | 0.003227 | 0.02729603 | 0.01000793 | 0.006141 | -0.001898739 | 0.005626773 | 0.7353 | -0.4451804 | 0.3473743 | 0.1921 | -0.1975919 | 0.2786004 | 0.4809 |
| 0.1 | 121 | CAF + TP + PX | -1.274 | 0.3532336 | 0.0006571 | 0.0298907 | 0.01470864 | 0.03303 | -0.001529166 | 0.007725097 | 0.86 | -0.1889974 | 0.5272425 | 0.7243 | -0.07701537 | 0.39117402 | 0.853 |
| 5 | 509 | CAF + TP + PX + MX | -0.823 | 0.2680991 | 0.002185 | 0.007905393 | 0.011399421 | 0.4884 | -0.001865078 | 0.006759327 | 0.7805 | -0.08618605 | 0.37925441 | 0.8215 | 0.06192471 | 0.32195188 | 0.8439 |
| 2 | 420 | CAF + TP + PX + MX | -1.03 | 0.2794238 | 0.0002277 | 0.02432609 | 0.01231021 | 0.04634 | -0.008733971 | 0.007285852 | 0.2239 | -0.1354184 | 0.4227936 | 0.7471 | -0.2425322 | 0.3602748 | 0.4988 |
| 1 | 276 | CAF + TP + PX + MX | -1.04 | 0.3276549 | 0.001572 | 0.01576093 | 0.01482524 | 0.2866 | -0.003045039 | 0.008400915 | 0.7095 | -0.01868127 | 0.51170186 | 0.9708 | -0.1721211 | 0.4128901 | 0.6739 |
| 0.1 | 121 | CAF + TP + PX + MX | -1.08 | 0.4919467 | 0.02253 | 0.01183208 | 0.02050152 | 0.5457 | 0.01806164 | 0.01014006 | 0.0739 | 0.1613058 | 0.7467483 | 0.802 | -0.3171138 | 0.5498856 | 0.5286 |
